# Changing the Motivation for Mental Work with Temporal Interference Stimulation

**DOI:** 10.1101/2025.09.24.678252

**Authors:** Gizem Vural, Sarina Drexler, Daniel Keeser, Alexander Soutschek

## Abstract

Successful goal-directed behavior often requires the exertion of effortful control processes. The striatum is thought to play a key role in determining whether a goal is worth the required cognitive effort, but whether and how the striatum causally influences effort-based decisions in humans remains unclear. Here, we address this gap by employing transcranial temporal interference stimulation (tTIS), a novel non-invasive brain stimulation technique capable of targeting deeper brain regions. We applied striatum-targeted tTIS to healthy participants performing an effort-based decision task during functional MRI scanning. Supporting the idea that the striatum encodes both the benefits and costs of actions, we found that, on the behavioral level, striatum-targeted tTIS increased the sensitivity to both reward magnitudes and effort costs. At the neural level, this was mirrored by stronger representations of effort demands in the striatum under stimulation as well as by enhanced functional coupling between the striatum and the anterior cingulate cortex, a region at the intersection of motivation and cognition. Together, our findings provide insights into the causal contributions of the striatum to trading-off rewards against effort costs, informing neural accounts of motivated cognition and suggesting novel neural interventions for the treatment of amotivation.

## Introduction

From learning for an exam to writing scientific articles, successful goal-directed behavior often requires cognitive control processes. However, people often avoid mentally demanding activities despite their beneficial consequences (Kool & Botvinick, 2014, 2018), suggesting that they trade off the benefits against the costs of effortful cognition. Moreover, pathologically reduced motivation for mental effort is a core symptom of several clinical conditions, including major depressive disorder, schizophrenia, and Parkinson’s Disease (Barch et al., 2023; Culbreth et al., 2016; Wolpe et al., 2024), and may contribute to the cognitive impairments in these disorders (Cooper et al., 2019; Moritz et al., 2017). For both basic and clinical research, it is therefore important to obtain a better understanding of the neural mechanisms underlying the motivation for mental effort.

Deciding whether a goal is worth the required effort costs is thought to be implemented by a distributed neural network in which the striatum, as part of the dopaminergic reward system, plays a key role (Bartra et al., 2013; Botvinick et al., 2009; Gershman et al., 2024; Salamone et al., 2007; Westbrook et al., 2019). The striatum encodes both reward magnitudes and the costs of actions, including cognitive effort (Westbrook et al., 2019; Westbrook et al., 2020), in the direct and indirect dopaminergic pathways, respectively. Recent models propose that the relative balance of the direct and indirect striatal pathway implements a cost-control signal that regulates the perceived affordability of mental effort relative to the value of the reward at stake (Beeler & Mourra, 2018; Collins & Frank, 2014; Soutschek et al., 2023). Activation in the direct and indirect pathway in turn influences neural processing of cost-benefit trade-offs in prefrontal regions such as the dorsal anterior cingulate cortex (ACC) and dorsolateral prefrontal cortex (dlPFC), which evaluate control demands and integrate cost-benefit information (Shenhav et al., 2013; Staudinger et al., 2011). The ACC in particular was ascribed a central role in deciding whether or not to engage in effortful control processes (implemented by the dlPFC) for goal achievement (Soutschek, Nadporozhskaia, et al., 2022; Walton et al., 2009). Via its structural connections with the striatum, the ACC may trade off reward against cost signals provided from the striatum to decide about control allocation (Croxson et al., 2009; Hauber & Sommer, 2009; Klein-Flugge et al., 2016; Silvestrini et al., 2022).

Despite the hypothesized role of the striatum for motivating mental effort, however, we have only a limited understanding of how activation in the striatum causally influences decisions to engage in goal-directed mental effort. This is because conventional non-invasive stimulation techniques, such as transcranial magnetic stimulation (TMS) or transcranial direct current stimulation (tDCS), allow manipulating activation only in cortical but not in subcortical regions like the striatum. Pharmacological interventions targeting the dopaminergic system, on the other hand, show systemic effects on dopamine receptors in the entire brain rather than selectively in the striatum. While animal studies provided evidence for the striatum’s causal involvement in effort-based decisions (Hauber & Sommer, 2009; Walton et al., 2009; Yohn et al., 2016), we therefore have a fundamental knowledge gap regarding how the striatum causally contributes to motivated mental effort in humans. This hampers both scientific theorizing and the development of treatments against amotivation in clinical disorders.

A potential solution to the problem of non-invasively modulating subcortical activation in humans is provided by transcranial temporal interference stimulation (tTIS) (Grossman et al., 2017). tTIS is based on the induction of two electric high-frequency fields beyond naturally occurring neural oscillation frequencies. In brain regions where the fields overlap, the interference of the high-frequency fields produces an envelope frequency that corresponds to the difference between the two high-frequency oscillations and modulates neural oscillations in these regions. It was recently shown that tTIS allows manipulating neural activation in subcortical areas like the hippocampus and striatum (Vassiliadis et al., 2024; Violante et al., 2023; Wessel et al., 2023). The goal of the current study, therefore, was to assess the causal role of the striatum for decisions to engage in mental effort by combining striatum-targeted tTIS with functional imaging. Based on the assumption that the striatum performs a cost control by computing the value of a reward relative to the associated costs (Beeler & Mourra, 2018), we hypothesized striatum-targeted tTIS to enhance the sensitivity to the benefits and costs of mental work in the striatum and connected areas such as ACC. This would provide a causal account of how the brain trades-off the value of goals against mental effort costs.

To test our hypothesis, participants made decisions whether or not to engage in a mentally effortful working memory task for monetary rewards under striatum-targeted versus sham tTIS in two separate experiments (Figure 1A/B). To obtain the rewards, participants had to perform different difficulty levels of the N-back task (0- to 4-back), which required them to maintain and update N letters in their working memory (Soutschek & Tobler, 2020; Westbrook et al., 2013; Westbrook et al., 2020). The goal of a first behavioral pilot experiment (11 participants) was to determine the optimal stimulation parameters. In a second experiment (33 participants), we then combined striatum-targeted tTIS with functional brain imaging (fMRI) to test the effects of tTIS on both the behavioral and the neural level. We hypothesized that, behaviorally, striatum-targeted tTIS would result in a stricter cost control with an increased sensitivity to both the anticipated benefits and the costs of mental effort. On the neural level, this should be mirrored by enhanced reward and cost signals in the striatum and stronger functional coupling between the striatum and cortical areas computing cost-benefit trade-offs like the ACC. Consistent with our hypotheses, our findings suggest a contribution of striatal activation to trading-off rewards against mental effort costs via communication with the ACC, providing insights into the brain networks causally underlying the motivation for goal-directed cognitive control.

**Figure 1.**
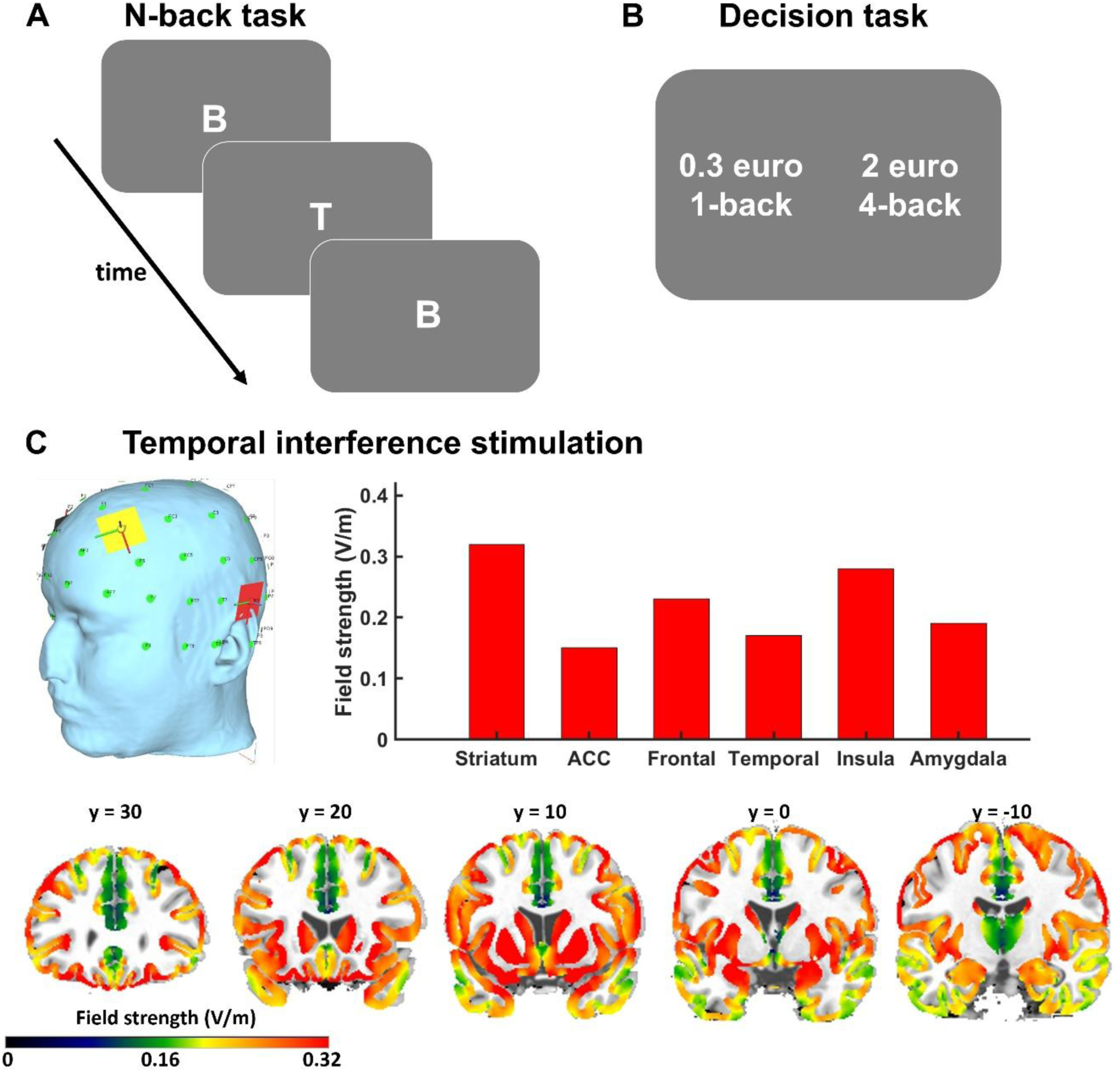
Experimental procedures. (A) To manipulate mental effort demands, participants performed varying levels of the N-back task (1- to 4-back) where they had to decide whether a currently presented letter was identical to the letter presented N trials before. (B) In the effort-based decision task (performed under tTIS in the MRI scanner), participants then made choices between low-effort (e.g., 0.3 euro for 1-back) and high-effort options (e.g., 2 euro for 4-back). (C) tTIS protocol: To target the striatum, we placed two electrode pairs over the bilateral prefrontal cortex and temporal lobes. Current flow simulations with SimNIBS suggest that the strength of the interference field peaked in the striatum and was lower in other regions involved in effort-based choice (ACC, insula, amygdala) or in the frontal and temporal regions where the electrodes were placed.

## Results

### Striatum-targeted tTIS enhances the sensitivity to the benefits and costs of cognition

Before examining the behavioral and neural effects of tTIS, we first conducted electric field simulations to confirm that our stimulation montage effectively targeted the striatum. Consistent with prior findings using a similar electrode configuration (Vassiliadis et al., 2024; Wessel et al., 2023), the simulations revealed peak electric field strengths within an anatomical mask of the striatum (combining nucleus accumbens, putamen and caudate: 0.33 V/m). The interference field was weaker in regions beneath the stimulation electrodes (bilateral dorsolateral prefrontal cortex: 0.27 V/m; bilateral middle temporal gyrus: 0.25 V/m) and areas involved in effort-based choice, including the ACC (0.16 V/m), insula (0.26 V/m), and amygdala (0.19 V/m; Figure 1C). These results suggest that the electrode configuration allows modulation of activation in subcortical areas, in particular the striatum where the electrical field was almost twice as strong compared to the ACC.

Next, we aimed to determine the optimal envelope frequency for changing cost-benefit computations with striatum-targeted tTIS. In a behavioral pilot experiment, we compared the effects of 6 Hz theta and 80 Hz gamma tTIS, relative to sham stimulation, on effort-based decisions. As expected, participants more strongly preferred high-effort options with increasing differences in reward magnitudes, β = 1.47, *z* = 3.02, *p* = 0.002, and decreasing differences in effort demands, β = –0.98, *z* = 2.63, *p* = 0.009, confirming that the task successfully elicited cost–benefit trade-offs. Importantly, we observed significant interactions between theta tTIS and differences in reward magnitudes, β = 1.71, *z* = 3.27, *p* = 0.001, as well as effort costs, β = −0.72, *z* = 1.97, *p* = 0.049, suggesting that theta tTIS enhanced the sensitivity to both rewards and effort demands during effort-based decisions (Figure 2A and Table 1). In contrast, we observed no significant impact of gamma tTIS on reward, β = 0.61, *z* = 1.16, *p* = 0.244, or effort processing, β = −0.33, *z* = 0.86, *p* = 0.392. These pilot results suggest that striatum-targeted theta, but not gamma, tTIS modulates cost-benefit decision making, providing an empirical rationale for selecting theta-frequency tTIS in the main study. The lack of significant gamma tTIS effects moreover suggests that the observed tTIS effects are specific to the envelope frequency and cannot be explained by direct stimulation effects of the carrier frequencies.

**Figure 2.**
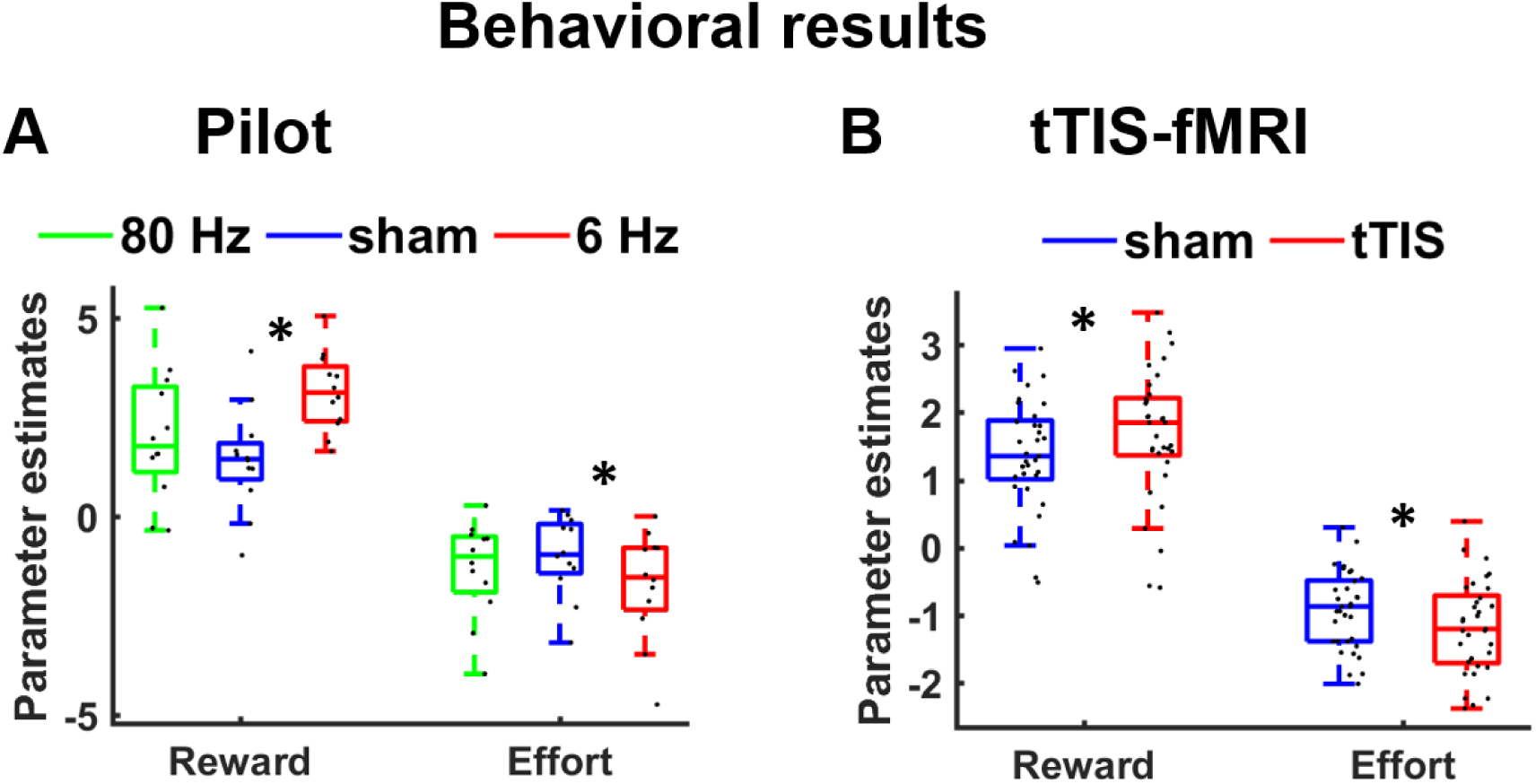
Impact of tTIS on choice behavior. (A) In the pilot experiment (outside the scanner), theta versus sham tTIS enhanced the sensitivity to both reward magnitudes and effort costs. Gamma tTIS, in contrast, showed no significant effects on behavior. (B) We replicated these findings in the main experiment inside the scanner: again, striatum-targeted theta tTIS strengthened the weights assigned to rewards and effort demands, consistent with the view that the striatum encodes both the benefits and costs of actions. For illustration purposes, we show individual regression weights from the hierarchical regression analyses. Asterisks indicate significant effects (* *p* < 0.05).

**Table 1.** Full mixed-effects regression results for the behavioral pilot experiment. The generalized linear mixed-effects model regressed binary choices (1 = high effort, 0 = low effort) on differences in reward magnitudes (Reward_diff_) and effort demands (Effort_diff_), tTIS frequency (6 Hz vs. sham, 80 Hz vs. sham), and all interaction effects. Discomfort and flicker ratings were included as control regressors. All continuous variables were z-standardized. *p < .05, **p < .01, ***p < .001.

| Fixed Effect | Estimate ($\beta$) | Std. Error | z value | p value |
| --- | --- | --- | --- | --- |
| Intercept | 1.66628 | 0.72889 | 2.286 | 0.0223* |
| 6 Hz tTIS | -0.29025 | 0.62079 | 0.468 | 0.6401 |
| 80 Hz tTIS | -0.05624 | 0.44903 | 0.125 | 0.9003 |
| $\text{Reward}_{\text{diff}}$ | 1.47275 | 0.48782 | 3.019 | 0.0025** |
| $\text{Effort}_{\text{diff}}$ | -0.98328 | 0.37462 | 2.625 | 0.0087** |
| Discomfort | -5.87236 | 2.19770 | 2.672 | 0.0075** |
| Flicker | 5.71636 | 2.19635 | 2.603 | 0.0093** |
| 6 Hz tTIS × $\text{Reward}_{\text{diff}}$ | 1.71306 | 0.52416 | 3.268 | 0.0011** |
| 80 Hz tTIS × $\text{Reward}_{\text{diff}}$ | 0.60633 | 0.52134 | 1.163 | 0.2448 |
| 6 Hz tTIS × $\text{Effort}_{\text{diff}}$ | -0.72748 | 0.36971 | 1.968 | 0.0491* |
| 80 Hz tTIS × $\text{Effort}_{\text{diff}}$ | -0.32883 | 0.38434 | 0.856 | 0.3922 |
| $\text{Reward}_{\text{diff}} \times \text{Effort}_{\text{diff}}$ | 0.74781 | 0.29383 | 2.545 | 0.0109* |
| 6 Hz tTIS × $\text{Reward}_{\text{diff}} \times \text{Effort}_{\text{diff}}$ | 0.75341 | 0.51635 | 1.459 | 0.1445 |
| 80 Hz tTIS × $\text{Reward}_{\text{diff}} \times \text{Effort}_{\text{diff}}$ | -0.57193 | 0.52104 | 1.098 | 0.2724 |

In the main experiment, 33 participants then performed the same effort-based decision task inside the MRI scanner while receiving theta versus sham tTIS targeting the striatum. Replicating the pilot findings, participants were more likely to choose high-effort options with increasing reward differences, β = 1.41, *z* = 7.78, *p* < 0.001, and decreasing differences in effort demands, β = −0.94, *z* = 6.57, *p* < 0.001. Importantly, theta tTIS again strengthened the sensitivity to both rewards, β = 0.31, z = 2.39, *p* = 0.017, and effort costs, β = −0.26, z = 2.08, *p* = 0.037 (Figure 2B and Table 2). Thus, two independent data sets provide converging evidence that striatum-targeted theta tTIS amplifies both cost and benefit considerations during effort-based decision-making, consistent with dual pathway accounts of the striatum’s role in motivating effortful cognition.

**Table 2.** Full mixed-effects regression results for the main experiment (inside the MRI scanner). The generalized linear mixed-effects model regressed binary choices (1 = high effort, 0 = low effort) on differences in reward magnitudes (Reward_diff_) and effort demands (Effort_diff_), tTIS frequency (6 Hz vs. sham), and all interaction effects. Discomfort and flicker ratings were included as control regressors. All continuous variables were z-standardized. *p < .05, **p < .01, ***p < .001.

| Fixed Effect | Estimate ($\beta$) | Std. Error | z value | p value |
| --- | --- | --- | --- | --- |
| Intercept | 0.91875 | 0.44436 | 2.068 | 0.0387* |
| 6 Hz tTIS | -0.14606 | 0.13679 | 1.068 | 0.2856 |
| $\text{Reward}_{\text{diff}}$ | 1.41285 | 0.18168 | 7.777 | <0.001*** |
| $\text{Effort}_{\text{diff}}$ | -0.94216 | 0.14349 | 6.566 | <0.001*** |
| Discomfort | 0.45674 | 0.29354 | 1.556 | 0.1197 |
| Flicker | 0.18666 | 0.26682 | 0.700 | 0.4842 |
| 6 Hz tTIS × $\text{Reward}_{\text{diff}}$ | 0.30816 | 0.12888 | 2.391 | 0.0168* |
| 6 Hz tTIS × $\text{Effort}_{\text{diff}}$ | -0.25898 | 0.12443 | 2.081 | 0.0374* |
| Reward diff × $\text{Effort}_{\text{diff}}$ | 0.18295 | 0.10193 | 1.795 | 0.0727. |
| 6 Hz tTIS × $\text{Reward}_{\text{diff}} \times \text{Effort}_{\text{diff}}$ | -0.05959 | 0.12529 | 0.476 | 0.6344 |

### Striatum-targeted tTIS strengthens effort-related activation in the striatum

Motivated by our behavioral results, we tested our neural hypothesis that striatum-targeted tTIS alters the neural encoding of reward and effort costs in the striatum, using a pre-defined region-of-interest (ROI) for the striatum based on meta-analyses of neural value coding (Bartra et al., 2013; Clithero & Rangel, 2014). Theta versus sham tTIS significantly increased the striatum’s sensitivity to effort costs, *t*(32) = 3.20, *p* = 0.003, Cohen’s *d* = 0.56 (Figure 3A/B): under theta tTIS, increasing effort demands were associated with a significant decrease in striatal activation, *t*(32) = 4.25, *p* < 0.001, Cohen’s *d* = 0.74, while we observed no significant effort-related activation under sham, *t*(32) = 1.01, *p* = 0.32, Cohen’s *d* = 0.18. The impact of theta versus sham tTIS on effort processing was robust to employing an anatomical striatum ROI, *t*(32) = 3.01, *p* = 0.005, Cohen’s *d* = 0.52. These findings are consistent with theoretical accounts proposing that the striatum encodes the expected costs of actions (Botvinick et al., 2009; Westbrook et al., 2019) and suggest that striatum-targeted tTIS enhances the striatum’s sensitivity to effort costs.

**Figure 3.**
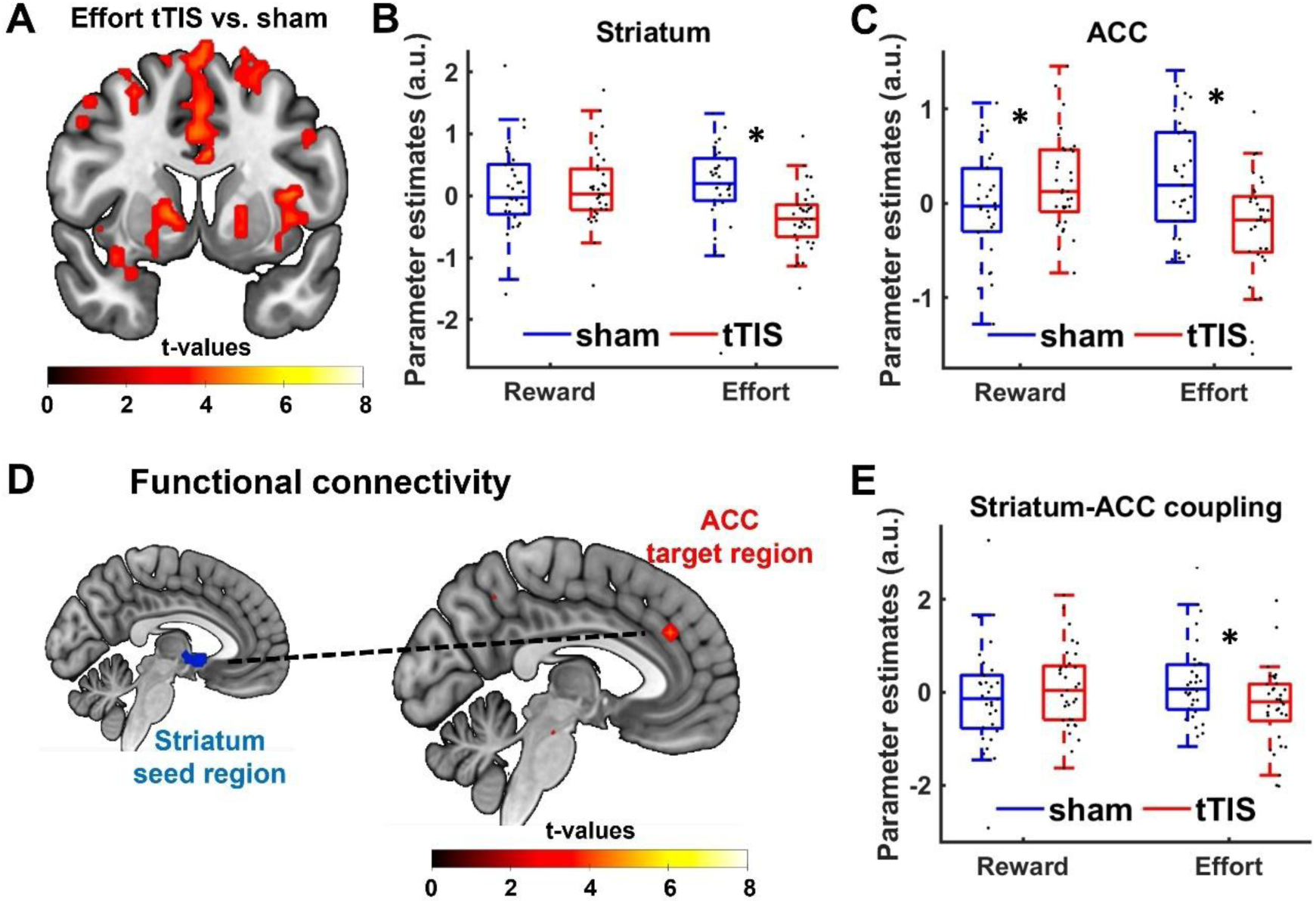
Effects of striatum-targeted tTIS on neural activation and functional connectivity. (A) Whole-brain activation map showing increased effort-related activity under active tTIS compared to sham. (B-C) Parameter estimates for reward and effort in the striatum and ACC, showing significant modulation of effort-related activity under tTIS. (D) PPI analysis using the striatum as a seed revealed increased connectivity with the ACC during effort-based decisions. (E) Striatum-ACC coupling estimates show significantly stronger connectivity under tTIS during effort-related decision-making. Error bars represent standard error of the mean. Asterisks indicate significant effects (* *p* < 0.05).

When we tested whether theta-frequency tTIS also modulated reward-related activation in the striatum, we found no evidence that striatal activation significantly correlated with differences in reward magnitude under sham, *t*(32) = 0.39, *p* = 0.70, Cohen’s *d* = 0.07, nor did striatal reward representations differ between stimulation conditions, *t*(32) = 0.80, *p* = 0.44, Cohen’s *d* = 0.14. One explanation for the absence of a reward effect in the imaging analysis is that reward levels in our task were individually adjusted based on participants’ indifference points for each effort level (see Materials and Methods for details), reducing variance in the differences between reward magnitudes and making our design more sensitive for detecting effort compared to reward signals in the striatum. In fact, when we computed a further GLM where the effort regressor was orthogonalized with respect to reward (such that the shared variance was assigned to the regressor for reward differences), we observed significant reward-related activation in the striatum ROI, *t*(32) = 2.20, *p* = 0.04, Cohen’s *d* = 0.38. This GLM also revealed a trend for tTIS versus sham effects on striatal reward representations, *t*(32) = 2.00, *p* = 0.06, Cohen’s *d* = 0.35, and replicated the stimulation effect on effort coding in the striatum, *t*(32) = 2.90, *p* = 0.007, Cohen’s *d* = 0.50, underlining the robustness of this effect. Lastly, exploratory whole-brain analyses revealed significant stimulation effects on effort representations in several regions including the bilateral supplementary motor areas (SMA), left primary somatosensory cortex (S1), and right middle temporal gyrus, all *p* < 0.05 family-wise error (FWE) corrected at cluster level (Table 3). For reward-related activation, no significant clusters were identified, all *p* > 0.70, FWE-corrected at cluster level. Taken together, our imaging results suggest that theta tTIS strengthens the representation of effort costs in the striatum.

**Table 3.**
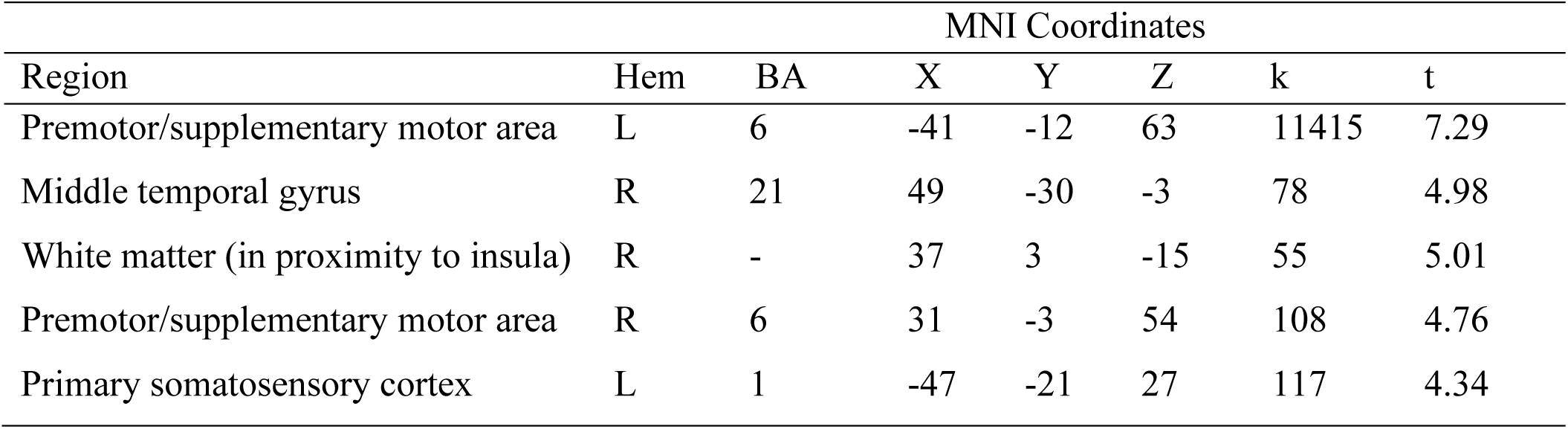
Results from the whole-brain contrast comparing effort-related activation during theta tTIS and sham stimulation. We report significant effects surviving family-wise error (FWE) correction at the cluster level (*p* < 0.05 with a cluster-inducing threshold of *p* < 0.001 uncorrected). Hem = Hemisphere (L = left, R = right); BA = Brodmann area

### tTIS modulates effort-dependent ACC-striatum coupling

Neural accounts of effort-based decisions assume that the striatum performs cost-benefit calculations not in isolation but in interaction with a prefrontal network, in particular the ACC (Shenhav et al., 2013; Silvestrini et al., 2022). In fact, when we investigated whether our stimulation affected also neural processing in the ACC ROI, we observed that theta versus sham tTIS strengthened both the representation of reward magnitudes, *t*(32) = 2.29, *p* = 0.029, Cohen’s *d* = 0.40, and effort costs *t*(32) = 3.51, *p* = 0.001, Cohen’s *d* = 0.61, the latter mirroring the findings in the striatum (Figure 3C). Note again that the electrical field was more than 90% stronger in the striatum than in the ACC, which suggests that the tTIS effects on the ACC are more likely to stem from a potential coupling with the striatum rather than from local stimulation effects. To assess whether striatum-targeted tTIS changed the communication between striatum and ACC, we performed a psycho-physiological interaction (PPI) analysis with the striatum ROI as seed region and tested for effort-related activation in the ACC ROI. Theta versus sham tTIS significantly affected effort-related striatum-ACC connectivity, *t*(32) = 2.32, *p* = 0.027, Cohen’s *d* = 0.40 (Figure 3D): While we observed no significant effort-related connectivity under sham, *t*(32) = 1.33, *p* = 0.19, Cohen’s *d* = 0.23, under theta tTIS we observed a trend for enhanced striatum-ACC coupling for decreasing differences in effort demands, *t*(32) = 1.95, *p* = 0.06, Cohen’s *d* = 0.34. This suggests that striatum-targeted tTIS changes the connectivity between striatum and ACC as a function of effort demands.

Exploratory whole-brain analyses revealed stronger coupling between striatum and frontopolar cortex for decreasing reward differences under sham, MNI peak coordinates: x = −32, y = 42, z = 15, *p* = 0.001 FWE-corrected at cluster level. This is consistent with the FPC’s role for motivating decisions to engage in rewarded mental effort (Bogdanov et al., 2025; Soutschek, Bagaini, et al., 2022; Soutschek et al., 2018). We observed no further clusters showing reward- or effort-related functional coupling with the striatum under sham or for theta versus sham tTIS, all *p* > 0.17 FWE-corrected at cluster level.

### Frontostriatal coupling mediates stimulation effects on choice behavior

The neural findings reported above raise the question as to whether the behavioral stimulation effects on reward and effort sensitivity can be explained by the influence of tTIS on the neural level. In the sham baseline condition, the strength of effort-related striatum-ACC coupling (extracted from the ACC ROI) correlated with the extracted individual coefficients for reward sensitivity, r = 0.36, *p* = 0.038, and effort sensitivity, r = −0.37, *p* = 0.033, from the behavioral regression analysis (Figure 4A/B). This suggests that stronger frontostriatal coupling for decreasing effort differences is associated with a stronger weighting of effort demands and a weaker reliance on reward differences in behavior. In contrast, activation in the striatum or ACC per se showed no correlations with reward or effort sensitivity from the behavioral analysis, all r < 0.16, all *p* > 0.37. This suggests that the striatum influences choice behavior via its coupling with the ACC rather than in isolation. To formally assess whether striatum-targeted tTIS affected behavior through changing striatum-ACC connectivity, we conducted a mediation analysis by adding individual coefficients for effort-related striatum-ACC connectivity to the behavioral regression analysis. While the influence of tTIS on reward sensitivity remained significant, β = 0.30, *z* = 2.15, *p* = 0.03, the stimulation effects on effort sensitivity showed only a non-significant trend-level effect, β = –0.27, *z* = 1.93, *p* = 0.05. Importantly, striatum-ACC coupling significantly predicted effort-based decisions, β *=* 0.46, *z* = 2.52, *p* = 0.01, and a Sobel test provided evidence for the significance of the indirect mediation path, *z* = 1.71, *p* = 0.04 (Figure 4C). This suggests that the impact of striatum-targeted tTIS on effort preferences can at least partially be explained by stimulation-induced changes in striatum-ACC connectivity.

**Figure 4.**
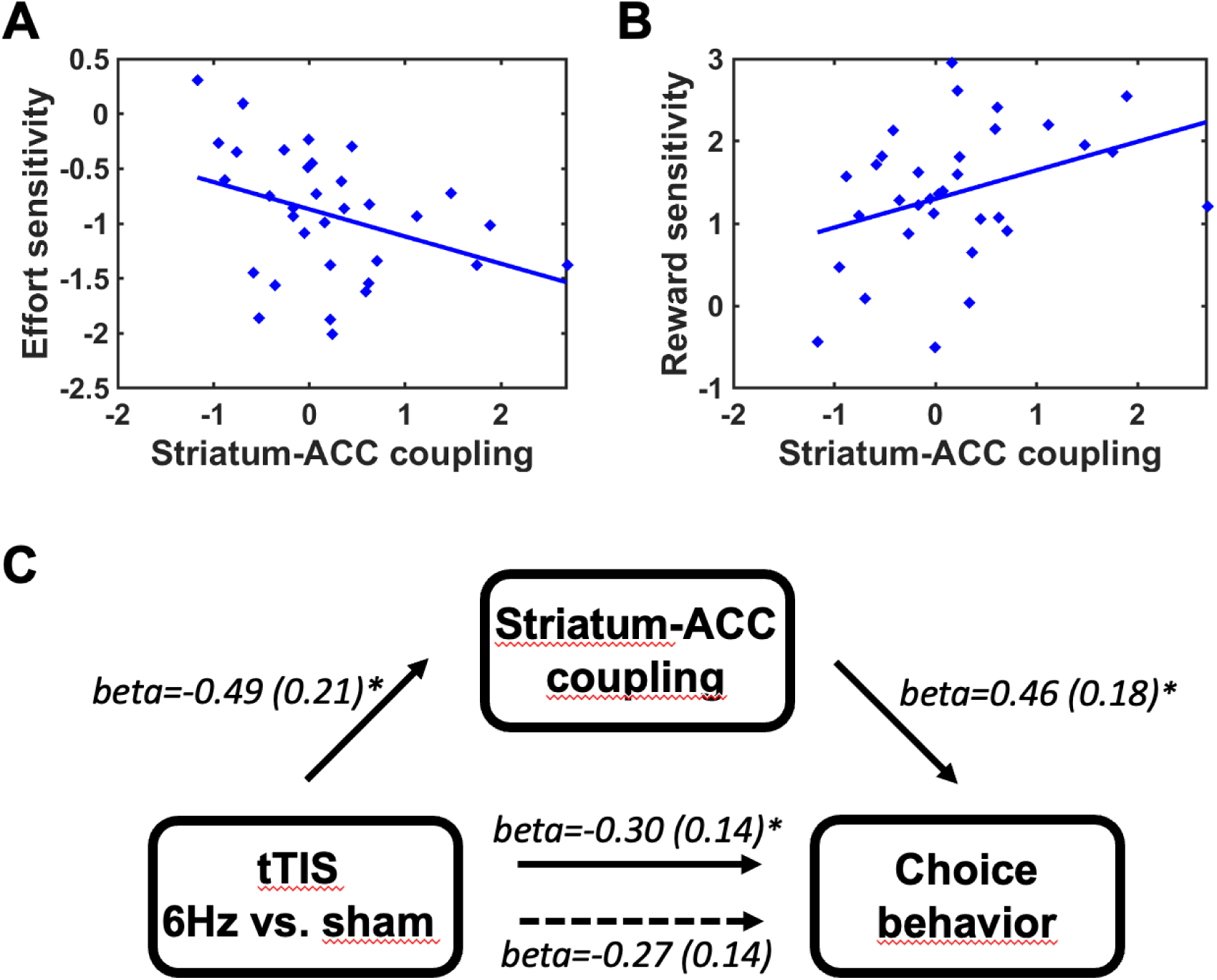
Brain-behavior mediation analysis. Under sham, effort-related striatum-ACC coupling was significantly correlated with individual differences in (A) effort and (B) reward sensitivity. (C) A mediation analysis (Sobel test) revealed a significant indirect mediation path: tTIS affected striatum-ACC coupling, which in turn influenced effort-based decisions. Because the impact of tTIS on effort sensitivity was no longer significant (though showed a trend-level effect) when controlling for striatum-ACC coupling (dashed line), we can explain the impact of tTIS on behavior at least partially via the stimulation effects on striatum-ACC coupling.

## Discussion

Decisions to engage in goal-directed mental effort are thought to rely on interactions between the striatum and the ACC (Hauber & Sommer, 2009; Silvestrini et al., 2022), but the striatum’s causal influence on these decisions remained unclear because conventional brain stimulation cannot manipulate deep subcortical structures. Here, we combined striatum-targeted tTIS with neuroimaging to determine how strengthening striatal theta oscillations affects behavioral and neural signatures of cost-benefit trade-offs. Behaviorally, the stimulation enhanced sensitivity to both reward magnitudes and effort demands during decision making. This aligns with dual-pathway models of the striatum, which propose that the direct pathway encodes the benefits of actions while the indirect pathway encodes the corresponding costs (Beeler & Mourra, 2018; Collins & Frank, 2014). Noteworthy, the stimulation did not affect overall preferences for high-effort choices, at variance with the assumption that striatal activation generally boosts the motivation for costly rewards (Yohn et al., 2016). On the neural level, these behavioral findings were mirrored by enhanced representation of effort costs in the striatum as well as stronger effort-related connectivity between the striatum and the ACC. Interestingly, it was the effort-dependent striatum-ACC coupling, rather than activation in the striatum per se, that correlated with individual differences in reward and effort sensitivity and that mediated stimulation effects on effort preferences on the behavioral level. This suggests that striatal representations of benefits and mental demands may not influence effort-based decisions directly but via their communication with the ACC, a region known for its role at the intersection of motivation and cognition (Holroyd & Yeung, 2012; Shenhav et al., 2013; Vriens et al., 2025). Together, our findings provide evidence that cost-benefit decision processes in humans can be altered through non-invasive stimulation targeting the striatum.

While several theoretical accounts ascribe the striatum a central role for cost-benefit trade-offs (Schmidt et al., 2012; Schouppe et al., 2014; Westbrook et al., 2019), evidence in humans so far has relied mainly on correlative data because most existing non-invasive intervention techniques do not allow manipulating subcortical activation. Our findings suggest that the striatum does not generally boost the motivation for costly rewards but strengthens the sensitivity to mental demands in effort-based decisions. This supports theoretical accounts according to which the striatum performs a “cost control” (Beeler & Mourra, 2018; Soutschek et al., 2023): The indirect “no go” pathway exerts inhibitory effects on cortical activation, preventing decision makers to engage in costly activities such as demanding mental work. High dopamine levels are thought to motivate goal-directed behavior by releasing the inhibitory effects of D2 receptors in the indirect pathway on cortical activation (Hikida et al., 2010; Zalocusky et al., 2016). Within this framework, we speculate that striatum-targeted tTIS strengthened particularly the inhibitory indirect pathway. This resulted in a stricter “cost control”, making individuals more sensitive to both the rewards at stake and the mental work required to obtain them. Therefore, entraining striatal theta oscillations with tTIS may produce effects distinct from enhanced dopamine levels, which were associated with a higher cost tolerance and thus a looser rather than stricter cost control (Beeler & Mourra, 2018; Soutschek et al., 2023; Webber et al., 2020). By supporting recent “cost control” accounts of the striatum, the current findings provide insights into the neural mechanisms underlying the striatum’s influence on goal-directed cognition.

Our results also suggest that the striatum motivates effortful cognition not in isolation but through its interaction with the ACC, a key region for higher-order control processes (Holroyd & Verguts, 2021; Holroyd & Yeung, 2012; Shenhav et al., 2013; Vriens et al., 2025). Theta tTIS increased functional coupling between ACC and striatum, consistent with the idea that communication between distant brain regions relies on synchronous theta oscillations (Canolty & Knight, 2010). The stimulation strengthened the representations of both rewards and costs in the ACC, supporting the idea that the ACC performs a cost-benefit analysis to decide whether or not to engage in goal-directed control processes (Shenhav et al., 2013; Vassena et al., 2014) and top-down influences action selection in the striatum if the expected benefits outweigh the costs (Fetcho et al., 2024). Theta oscillations in ACC were also related to decisions for mental effort and to higher-order control processes (Soutschek, Nadporozhskaia, et al., 2022; van Driel et al., 2015). Our results therefore provide further support for recent motivational perspectives on the ACC, underlining its key role at the intersection of motivation and cognition in concert with the striatum acting as a cost controller. As a caveat, we note that we cannot formally rule out direct stimulation effects on the ACC or other regions like insula or amygdala. However, the interference field in the striatum was considerably stronger than in these areas (+113%, +14%, and +68% than in ACC, insula, and amygdala, respectively). Moreover, our mediation analysis suggests that stimulation effects on the striatum’s connectivity with ACC at least partially explain the tTIS-induced behavioral changes, suggesting that changes in ACC activation alone are not sufficient to explain the tTIS effects on motivation. Lastly, past neurostimulation studies targeting the ACC reported a generally increased motivation for mental effort under stimulation, rather than increased benefit and cost sensitivity (Soutschek, Nadporozhskaia, et al., 2022). On balance, it seems thus more likely that the observed findings are to be explained by subcortical stimulation effects on the striatum rather than on cortical regions like the ACC.

On a general level, the suggested causal link between frontostriatal coupling and effort-based decisions informs debates about how the motivation for mental effort may change over the lifespan due to brain maturation (Galvan, 2014) or may be pathologically altered as a result of brain dysfunctions. Impairments in the motivation for goal-directed mental effort are a defining symptom of several clinical disorders (Wolpe et al., 2024) and may contribute to the deficits in higher-order cognitive functioning (Cooper et al., 2019; Moritz et al., 2017). By showing how the striatum influences cost-benefit computations, the current findings improve our mechanistic understanding of how striatal dysfunctions change motivational processes in these disorders. Moreover, abnormal activation in the striatum was related not only to amotivation but also to further symptoms in clinical disorders like impulsiveness, compulsivity, or motor dysfunctions (Harada et al., 2021; Kalivas & Volkow, 2005; Obeso et al., 2008). Pharmacological treatments of these symptoms lack regional specificity as they show systemic effects on neural activity, whereas established non-invasive brain stimulation approaches do not allow modulating subcortical activation directly. tTIS has the potential to overcome these limitations by providing a tool for manipulating subcortical activation. In the long run, this may stimulate the development of novel therapeutic neural interventions for clinical symptoms related to dysfunctions in subcortical brain networks.

To conclude, the motivation for goal-directed effortful cognition is key to academic achievements as well as professional success. Here, we identify a neural mechanism that causally underlies the motivation for mental work and show that this motivation can be changed by a novel brain stimulation approach targeting subcortical neural mechanisms. The current findings advance our understanding of the causal neural mechanisms underlying goal-directed mental effort and open new avenues for developing targeted neural interventions.

## Materials and Methods

### Participants

For the main experiment, we collected data from 33 participants (22 female, mean age: 22.45 years, standard deviation (SD): 3.71) who were recruited from the participant pool of the Department of Psychology at Ludwig Maximilian University (LMU) Munich. According to an a priori power analysis based on the effect size from a previous tACS study on value-based choice (Soutschek et al., 2021), a sample size of 30 participants should be sufficient to detect an effect with a power of 80% and an alpha level of 5%. A further sample of 11 volunteers (6 female, mean age: 24.92 years, SD: 2.18) participated in the pilot experiment. All participants provided informed written consent before participation. The study was approved by the local ethics committee of the psychology department at LMU Munich. Participants were screened for contraindications related to brain stimulation and MRI safety. They received a fixed compensation of 40 Euros for their participation, along with an additional performance-based bonus (see below).

### Stimuli and Task Design

The effort-based decision task was implemented using the Cogent 2000 toolbox in MATLAB. Participants repeatedly chose between two options differing in required cognitive effort and associated monetary reward (Figure 1A). Effort demands were manipulated using an N-back task ranging from 0-back (low effort) to 4-back (high effort), while reward magnitudes ranged from 0€ to 2€. Each trial displayed two options on the left and right sides of the screen, one high-effort option (1-back to 4-back level) and one low-effort option (0-back to 3-back). The task comprised 10 combinations of effort levels, each with 6 different reward levels. While the reward magnitude of the high-effort option was fixed to 2€, reward levels of the low-effort option were determined based on individual indifference points for the different effort combinations at the start of an experimental session using a titration approach. In more detail, in the calibration phase the reward for the low-effort option started at 1€ (high-effort reward was fixed to 2€) for each combination of low-effort and high-effort demands and was then adjusted by adding (after high-effort choices) or subtracting (after low-effort choices) 50% of the low-effort reward magnitude. Participants made two choices for each combination of effort levels in the calibration phase at the start of the experiment, and the resulting adjusted reward magnitudes of the low-effort option were considered as indifference points. For the main experiment (under tTIS), reward magnitudes of the low effort option were adjusted around each participant’s indifference points in increments of ±0.05, ±0.15, and ±0.25€. This was done to ensure that we measured participants’ effort preferences in their individual sensitive range, increasing the likelihood of observing significant stimulation effects on choice behavior. We used 8 levels for the low-effort reward (−0.25, −0.15, −0.05, −0.05, +0.05, +0.05, +0.15, and +0.25 around indifference point) for each of the 10 combinations of effort levels, resulting in a total of 80 trials.

On each trial, participants had 4 seconds to select one of the options via pressing a left or right response key (for the left or right choice option, respectively) on an MR-compatible button box. Each decision trial was followed by a Poisson-distributed jittered inter-trial interval (ITI; average = 3 seconds).

We informed participants that at the end of the experiment we would randomly select one of the choices they had made during the decision task, and that they would need to perform the selected N-back level for 2 minutes in order to obtain the monetary bonus. Following previous procedures (Soutschek, Bagaini, et al., 2022; Soutschek et al., 2018; Westbrook et al., 2020), mental effort had to be exerted at the end of the experiment rather than after each choice to disentangle effort preferences from fatigue effects.

### Procedure

The experiment consisted of two phases. In the first phase (outside the MRI scanner), participants completed a practice (calibration) session to familiarize themselves with the N-back levels and the decision task. The choices from the practice task were used to calibrate the reward levels of the decision task inside the scanner to the individual indifference points (see above).

In the second phase (inside the scanner), participants performed the calibrated version of the decision task under both active and sham stimulation (within-subject design). The order of stimulation conditions was counterbalanced across participants using four pseudorandomized sequences. Participants completed four runs, each consisting of two miniblocks. Every miniblock included 20 trials (160 trials in total; 80 trials per stimulation condition). At the end of each miniblock, participants rated their perceived discomfort (from 0 = “not painful at all” to 20 = “very painful”) and flicker perception (from 0 = “no flicker perceived” to 20 = “very strong flicker perception”) within 5 seconds. Note that discomfort and flicker ratings did not significantly differ between active and sham stimulation, discomfort: *t*(32) = 0.94, *p* = 0.35, flicker: *t*(32) = 1.40, *p* = 0.17, suggesting that blinding of tDCS conditions was successful. Short breaks of 30 seconds were included between miniblocks to minimize stimulation-induced aftereffects (Christian et al., 2023; Moisa et al., 2016). Longer breaks between runs were used for acquiring structural MRI scans (T1/T2-weighted), during which no stimulation or task was administered. The total fMRI session lasted approximately 45 minutes.

### tTIS Protocol

tTIS was administered using an MR-compatible, multi-channel DC-STIMULATOR MC (neuroConn GmbH, Ilmenau, Germany). In the main experiment, the stimulation protocol consisted of two conditions: active tTIS at a 6 Hz theta frequency and sham stimulation. In the pilot experiment, we additionally used an 80 Hz frequency.

Stimulation was delivered via two pairs of MR-compatible squared rubber electrodes (3 × 3 cm), placed at positions F3-F4 and TP7-TP8 according to the international 10-20 EEG system. We note that some previous tTIS study, but field simulations suggested that the induced interference fields show only marginal differences for 3 × 3 cm and high-definition (diameter = 2 cm) electrodes. Electrode placement was guided using a customized EEG cap with predefined markers to ensure consistent positioning across participants. Electrodes were affixed with Ten20 conductive paste (Weaver and Company) and secured with elastic bandages. The electrode configuration was chosen to target deep brain structures, particularly the basal ganglia, following the electrode setup in prior tTIS studies on motor learning (Vassiliadis et al., 2024; Wessel et al., 2023).

For the active stimulation condition, channel 1 delivered a 600 Hz current and channel 2 delivered 606 Hz, resulting in an interference pattern with a 6 Hz envelope frequency for the overlapping electric fields. For the sham condition, both channels delivered 600 Hz, producing no envelope frequency. In the pilot experiment, we additionally employed frequencies of 600 Hz and 680 Hz at channels 1 and 2, respectively, to induce an envelope frequency in the 80 Hz gamma range. The stimulation intensity was set to 1.8 mA (3.6 mA peak-to-peak). We employed carrier frequencies that were below the frequencies in some other tTIS studies (Vassiliadis et al., 2024; Violante et al., 2023; Wessel et al., 2023) because frequencies above 600 Hz might dampen the intensity of the stimulation (email communication with technical service from Neuroconn). Nevertheless, stimulation frequencies of 600 Hz are above the oscillation frequencies naturally occurring in the brain (Kucewicz et al., 2024), and also the lack of significant results in the 80 Hz tTIS condition (see Results section) speaks against the idea that the observed stimulation effects are driven by the carrier rather than the envelope frequencies. A ramp-up and ramp-down period of 5 seconds was applied at the beginning and end of each stimulation block. After the current ramp-up period, we stimulated participants for 25 seconds before the start of the decision task, allowing tTIS effect to build up prior to task performance. Stimulation during task performance lasted for 160 s, then the current was ramped down directly after the decision task. In the sham condition, we stimulated participants prior to task performance to simulate the tactile sensations of active stimulation, but the current was ramped down before the start of the task. Electrode impedance was monitored throughout the session and remained within acceptable limits for all participants.

To validate that our tTIS setup induced an interference field in the striatum, in line with previous work (Vassiliadis et al., 2024; Wessel et al., 2023), we performed current flow simulations using SimNIBS 4.1 (Saturnino et al., 2019). We first simulated the electrical fields separately for each electrode pair and then computed the interference field using the get_maxTI function. We assessed the field strength of the interference field with anatomical ROIs for the striatum, ACC, insula, amygdala, as well as the areas under the electrodes (bilateral DLPFC and middle temporal gyrus).

### MRI Data Acquisition and Preprocessing

MRI data were acquired using a 3 Tesla Siemens PRISMA scanner (Siemens Healthcare, Erlangen, Germany) with a 64-channel phased-array head coil. Functional images were collected using a T2*-weighted echo-planar imaging (EPI) sequence (TR = 1000 ms, TE = 30 ms, flip angle = 45°, 48 slices, voxel size = 3 × 3 × 3 mm, field of view = 210 × 240 mm, multiband factor = 4, 415 volumes per run). To improve whole-brain coverage and minimize susceptibility-related signal loss in ventral regions, slices were acquired in an oblique orientation tilted 3.3° toward the coronal plane.

Structural images included a high-resolution T1-weighted scan acquired using a 3D MPRAGE sequence (TR = 2500 ms, TE = 2.22 ms, flip angle = 8°, voxel size = 0.8 mm isotropic, 208 sagittal slices, FoV = 256 × 240 mm) and a T2-weighted 3D SPACE scan (TR = 5000 ms, TE = 383 ms, voxel size = 1 × 1 × 1 mm, 176 slices).

Functional data preprocessing was carried out in MATLAB using SPM12. The pipeline included the following steps: realignment and unwarping of functional volumes, slice timing correction, coregistration of the mean EPI image to the individual T1-weighted anatomical image, segmentation of the structural scan, normalization to MNI space with a resampling resolution of 3 x 3 x 3 mm, and spatial smoothing with a 6 mm full-width at half-maximum (FWHM) Gaussian kernel. A high-pass temporal filter with a cutoff of 128 seconds was applied to remove low-frequency signal drifts. Preprocessed images were visually inspected to confirm alignment and normalization quality. One participant was excluded from all analyses due to excessive head motion (> 6 mm).

### Statistical Analyses

#### Behavioral data

Statistical analyses of the behavioral data were performed using R (version 4.2.2). The significance level was set at alpha = 0.05 for all analyses. In the decision task, reaction times (RTs) faster than 250 ms were excluded from the analysis. Only one trial (86 ms) from the main experiment was removed due to this criterion.

For the analysis of binary choices in the pilot study (which aimed at determining the optimal stimulation frequency for the main study), we computed a generalized linear mixed-effects model (GLMM) using the lme4 package in R (Bates et al., 2014), assuming a binomial distribution with a logit link function. The GLMM regressed binary choices (0 = low-effort choice, 1 = high-effort choice) on fixed-effect predictors for tTIS (6 Hz vs. sham, 80 Hz vs. sham), differences in reward magnitude, differences in effort demands, and all interaction effects. To control for potential sensory confounds, discomfort and flicker ratings (averaged per participant and stimulation condition) were included as control variables of no interest. All continuous predictors were z-transformed. All fixed effects were also modelled as random slopes in addition to participant-specific random intercepts. The same GLMM was also used to analyze choice data in the main experiment, with the only difference that the main experiment did not include an 80 Hz tTIS condition.

#### Imaging analyses

To examine the effects of tTIS on neural processing, we computed first-level general linear models (GLMs) on the preprocessed functional data in SPM12. We modelled separate onset regressors for the sham and active stimulation conditions, which were time-locked to the offer presentation period during which participants evaluated the two choice options (duration = 4 seconds). To examine the neural correlates of reward and effort processing, z-transformed reward and effort differences were included as parametric modulators for each onset regressor (without orthogonalization). Six motion parameters from the realignment were included as nuisance regressors to control for head motion. Response omission trials were excluded. We computed individual contrast images for the main effects of reward and effort under sham, under tTIS, as well as for the difference between tTIS and sham.

To assess our neural hypotheses at the group level, we performed second-level analyses by entering individual contrast images from the first-level analysis into one-sample t-tests. For whole-brain analyses, statistical maps were thresholded with cluster-level family-wise error (FWE) correction at *p* < 0.05, using a cluster-inducing voxel-wise threshold of *p* < 0.001 (Eklund et al., 2016).

To examine our a priori hypotheses about striatal and ACC involvement in effort-based decision making, we defined region of interest (ROI) masks for these areas. For the striatum, we defined both an anatomical mask based on the AAL atlas in SPM12 (comprising bilateral putamen, caudate, and nucleus accumbens) and a functional mask based on published meta-analyses on neural value coding (Bartra et al., 2013; Clithero & Rangel, 2014). This mask can be downloaded from the following webpage: https://www.rnl.caltech.edu/resources/index.html. We defined two striatum ROIs to test the robustness of our findings. For the ACC ROI, we used an anatomical mask from the AAL atlas. We used these ROIs to test for tTIS on neural activation in the striatum and ACC, both via extracting parameter estimates from these ROIs with the MarsBar toolbox (Brett et al., 2002).

#### Psycho-physiological interaction (PPI) analysis

To test the hypothesis that tTIS changes the functional connectivity of the striatum with other regions involved in effort-based decision-making (such as the ACC), we performed a PPI analysis with the meta-analysis-based striatum ROI as seed region. We first extracted the time course of the BOLD signal from the striatum ROI as a physiological regressor and convolved it with psychological regressors for (i) reward and (ii) effort differences, separately for active and sham tTIS, to create PPI terms. The four PPI terms (reward and effort differences under tTIS and under sham) as well as the psychological and physiological regressors were then added to our original first-level GLM. We computed participant-specific contrasts for each PPI term in the first-level analysis and submitted them to second-level one-sample t-tests. To test our a priori hypothesis that tTIS changes fronto-striatal coupling, we used the anatomical ACC ROI to assess whether activation in this ROI is functionally coupled with the seed region in the striatum. We again conducted exploratory whole-brain analyses with cluster-level FWE correction (*p* < 0.05, cluster-inducing threshold of *p* < 0.001 uncorrected).

#### Brain-behavior mediation analysis

To examine whether the behavioral effects of theta-frequency tTIS were mediated by effort-related fronto-striatal connectivity, we conducted a mediation analysis. Specifically, we first tested whether theta versus sham tTIS modulated functional coupling between the striatum and the ACC, based on the extracted regression weights from the PPI analysis (path a). In the second step, we then added regression weights for striatum-ACC connectivity as a further predictor to our original GLMM on behavioral choices. This allowed us to assess whether striatum-ACC connectivity significantly predicted behavioral choices (path b) as well as whether the tTIS effects on behavior remained significant when controlling for tTIS-induced changes in striatum-ACC connectivity (path c’). To formally assess the significance of the indirect path, we conducted a one-tailed Sobel test using the unstandardized regression coefficients and standard errors from path a and path b to assess the directed hypothesis that the impact of tTIS on choices is mediated by changes in fronto-striatal connectivity.

## Acknowledgements

AS was supported by an Emmy Noether scholarship from the German Research Foundation (SO 1636/2-2) and an Exploration grant from the Boehringer Ingelheim Foundation. We are grateful to Türkay Sahin and Yagmur Karakoc for help with data collection.

## Data and code availability statement

The behavioral raw data underlying the results reported in this manuscript and the analysis code will be publicly available on the Open Science Framework (OSF; https://osf.io/5n9hs/overview?view_only=baa852d02c314783b3b804efa3f805f0). Functional imaging data will be made available upon reasonable request.

## Ethics statement

The study was approved by the local ethics committee of the psychology department at the LMU Munich.

## References

Barch, D. M., Culbreth, A. J., Ben Zeev, D., Campbell, A., Nepal, S., & Moran, E. K. (2023). Dissociation of Cognitive Effort-Based Decision Making and Its Associations With Symptoms, Cognition, and Everyday Life Function Across Schizophrenia, Bipolar Disorder, and Depression. Biol Psychiatry, 94(6), 501–510. 10.1016/j.biopsych.2023.04.007

Bartra, O., McGuire, J. T., & Kable, J. W. (2013). The valuation system: a coordinate-based meta-analysis of BOLD fMRI experiments examining neural correlates of subjective value. Neuroimage, 76, 412–427. 10.1016/j.neuroimage.2013.02.063

Bates, D., Mächler, M., Bolker, B., & Walker, S. (2014). Fitting linear mixed-effects models using lme4. arXiv preprint arXiv:1406.5823.

Beeler, J. A., & Mourra, D. (2018). To do or not to do: dopamine, affordability and the economics of opportunity. Frontiers in integrative neuroscience, 12, 6.

Bogdanov, M., Bustamante, L. A., Devine, S., Sheldon, S., & Otto, A. R. (2025). Noninvasive Brain Stimulation over the Frontopolar Cortex Promotes Willingness to Exert Cognitive Effort in a Foraging-Like Sequential Choice Task. J Neurosci, 45(10). 10.1523/JNEUROSCI.0647-24.2024

Botvinick, M. M., Huffstetler, S., & McGuire, J. T. (2009). Effort discounting in human nucleus accumbens. Cogn Affect Behav Neurosci, 9(1), 16–27. 10.3758/CABN.9.1.16

Brett, M., Anton, J.-L., Valabregue, R., & Poline, J.-B. (2002). Region of interest analysis using the MarsBar toolbox for SPM 99. Neuroimage, 16(2), S497.

Canolty, R. T., & Knight, R. T. (2010). The functional role of cross-frequency coupling. Trends Cogn Sci, 14(11), 506–515. 10.1016/j.tics.2010.09.001

Christian, P., Kapetaniou, G. E., & Soutschek, A. (2023). Causal roles of prefrontal and temporo-parietal theta oscillations for inequity aversion. Soc Cogn Affect Neurosci, nsad061. 10.1093/scan/nsad061

Clithero, J. A., & Rangel, A. (2014). Informatic parcellation of the network involved in the computation of subjective value. Soc Cogn Affect Neurosci, 9(9), 1289–1302. 10.1093/scan/nst106

Collins, A. G., & Frank, M. J. (2014). Opponent actor learning (OpAL): modeling interactive effects of striatal dopamine on reinforcement learning and choice incentive. Psychol Rev, 121(3), 337.

Cooper, J. A., Barch, D. M., Reddy, L. F., Horan, W. P., Green, M. F., & Treadway, M. T. (2019). Effortful goal-directed behavior in schizophrenia: Computational subtypes and associations with cognition. J Abnorm Psychol, 128(7), 710–722. 10.1037/abn0000443

Croxson, P. L., Walton, M. E., O’Reilly, J. X., Behrens, T. E., & Rushworth, M. F. (2009). Effort-based cost-benefit valuation and the human brain. J Neurosci, 29(14), 4531–4541. 10.1523/JNEUROSCI.4515-08.2009

Culbreth, A., Westbrook, A., & Barch, D. (2016). Negative symptoms are associated with an increased subjective cost of cognitive effort. J Abnorm Psychol, 125(4), 528–536. 10.1037/abn0000153

Eklund, A., Nichols, T. E., & Knutsson, H. (2016). Cluster failure: Why fMRI inferences for spatial extent have inflated false-positive rates. Proc Natl Acad Sci U S A, 113(28), 7900–7905. 10.1073/pnas.1602413113

Fetcho, R. N., Parekh, P. K., Chou, J., Kenwood, M., Chalençon, L., Estrin, D. J., … Liston, C. (2024). A stress-sensitive frontostriatal circuit supporting effortful reward-seeking behavior. Neuron, 112(3), 473–487. e474.

Galvan, A. (2014). Neural systems underlying reward and approach behaviors in childhood and adolescence. Curr Top Behav Neurosci, 16, 167–188. 10.1007/7854_2013_240

Gershman, S. J., Assad, J. A., Datta, S. R., Linderman, S. W., Sabatini, B. L., Uchida, N., & Wilbrecht, L. (2024). Explaining dopamine through prediction errors and beyond. Nat Neurosci, 27(9), 1645–1655.

Grossman, N., Bono, D., Dedic, N., Kodandaramaiah, S. B., Rudenko, A., Suk, H. J., … Boyden, E. S. (2017). Noninvasive Deep Brain Stimulation via Temporally Interfering Electric Fields. cell, 169(6), 1029–1041 e1016. 10.1016/j.cell.2017.05.024

Harada, M., Pascoli, V., Hiver, A., Flakowski, J., & Luscher, C. (2021). Corticostriatal Activity Driving Compulsive Reward Seeking. Biol Psychiatry, 90(12), 808–818. 10.1016/j.biopsych.2021.08.018

Hauber, W., & Sommer, S. (2009). Prefrontostriatal circuitry regulates effort-related decision making. Cerebral Cortex, 19(10), 2240–2247.

Hikida, T., Kimura, K., Wada, N., Funabiki, K., & Nakanishi, S. (2010). Distinct roles of synaptic transmission in direct and indirect striatal pathways to reward and aversive behavior. Neuron, 66(6), 896–907. 10.1016/j.neuron.2010.05.011

Holroyd, C. B., & Verguts, T. (2021). The Best Laid Plans: Computational Principles of Anterior Cingulate Cortex. Trends Cogn Sci, 25(4), 316–329. 10.1016/j.tics.2021.01.008

Holroyd, C. B., & Yeung, N. (2012). Motivation of extended behaviors by anterior cingulate cortex. Trends Cogn Sci, 16(2), 122–128. 10.1016/j.tics.2011.12.008 S1364-6613(11)00259-2 [pii]

Kalivas, P. W., & Volkow, N. D. (2005). The neural basis of addiction: a pathology of motivation and choice. Am J Psychiatry, 162(8), 1403–1413. 10.1176/appi.ajp.162.8.1403

Klein-Flugge, M. C., Kennerley, S. W., Friston, K., & Bestmann, S. (2016). Neural Signatures of Value Comparison in Human Cingulate Cortex during Decisions Requiring an Effort-Reward Trade-off. J Neurosci, 36(39), 10002–10015. 10.1523/JNEUROSCI.0292-16.2016

Kool, W., & Botvinick, M. (2014). A labor/leisure tradeoff in cognitive control. J Exp Psychol Gen, 143(1), 131–141. 10.1037/a0031048

Kool, W., & Botvinick, M. (2018). Mental labour. Nature Human Behaviour, 2(12), 899–908. 10.1038/s41562-018-0401-9

Kucewicz, M. T., Cimbalnik, J., Garcia-Salinas, J. S., Brazdil, M., & Worrell, G. A. (2024). High frequency oscillations in human memory and cognition: a neurophysiological substrate of engrams? Brain, 147(9), 2966–2982. 10.1093/brain/awae159

Moisa, M., Polania, R., Grueschow, M., & Ruff, C. C. (2016). Brain Network Mechanisms Underlying Motor Enhancement by Transcranial Entrainment of Gamma Oscillations. J Neurosci, 36(47), 12053–12065. 10.1523/JNEUROSCI.2044-16.2016

Moritz, S., Stockert, K., Hauschildt, M., Lill, H., Jelinek, L., Beblo, T., … Arlt, S. (2017). Are we exaggerating neuropsychological impairment in depression? Reopening a closed chapter. Expert Rev Neurother, 17(8), 839–846. 10.1080/14737175.2017.1347040

Obeso, J. A., Rodriguez-Oroz, M. C., Benitez-Temino, B., Blesa, F. J., Guridi, J., Marin, C., & Rodriguez, M. (2008). Functional organization of the basal ganglia: therapeutic implications for Parkinson’s disease. Mov Disord, 23 Suppl 3, S548–559. 10.1002/mds.22062

Salamone, J. D., Correa, M., Farrar, A., & Mingote, S. M. (2007). Effort-related functions of nucleus accumbens dopamine and associated forebrain circuits. Psychopharmacology (Berl), 191(3), 461–482. 10.1007/s00213-006-0668-9

Saturnino, G. B., Puonti, O., Nielsen, J. D., Antonenko, D., Madsen, K. H., & Thielscher, A. (2019). SimNIBS 2.1: a comprehensive pipeline for individualized electric field modelling for transcranial brain stimulation. Brain and human body modeling, 3–25.

Schmidt, L., Lebreton, M., Clery-Melin, M. L., Daunizeau, J., & Pessiglione, M. (2012). Neural mechanisms underlying motivation of mental versus physical effort. PLoS Biol, 10(2), e1001266. 10.1371/journal.pbio.1001266

Schouppe, N., Demanet, J., Boehler, C. N., Ridderinkhof, K. R., & Notebaert, W. (2014). The role of the striatum in effort-based decision-making in the absence of reward. J Neurosci, 34(6), 2148–2154. 10.1523/JNEUROSCI.1214-13.2014

Shenhav, A., Botvinick, M. M., & Cohen, J. D. (2013). The expected value of control: an integrative theory of anterior cingulate cortex function. Neuron, 79(2), 217–240. 10.1016/j.neuron.2013.07.007

Silvestrini, N., Musslick, S., Berry, A. S., & Vassena, E. (2022). An integrative effort: Bridging motivational intensity theory and recent neurocomputational and neuronal models of effort and control allocation. Psychol Rev.

Soutschek, A., Bagaini, A., Hare, T. A., & Tobler, P. N. (2022). Reconciling psychological and neuroscientific accounts of reduced motivation in aging. Soc Cogn Affect Neurosci, 17(4), 398–407. 10.1093/scan/nsab101

Soutschek, A., Jetter, A., & Tobler, P. N. (2023). Toward a Unifying Account of Dopamine’s Role in Cost-Benefit Decision Making. Biol Psychiatry Glob Open Sci, 3(2), 179–186. 10.1016/j.bpsgos.2022.02.010

Soutschek, A., Kang, P., Ruff, C. C., Hare, T. A., & Tobler, P. N. (2018). Brain Stimulation Over the Frontopolar Cortex Enhances Motivation to Exert Effort for Reward. Biol Psychiatry, 84(1), 38–45. 10.1016/j.biopsych.2017.11.007

Soutschek, A., Moisa, M., Ruff, C. C., & Tobler, P. N. (2021). Frontopolar theta oscillations link metacognition with prospective decision making. Nat Commun, 12(1), 3943. 10.1038/s41467-021-24197-3

Soutschek, A., Nadporozhskaia, L., & Christian, P. (2022). Brain stimulation over dorsomedial prefrontal cortex modulates effort-based decision making. Cogn Affect Behav Neurosci, 22(6), 1264–1274. 10.3758/s13415-022-01021-z

Soutschek, A., & Tobler, P. N. (2020). Causal role of lateral prefrontal cortex in mental effort and fatigue. Hum Brain Mapp, 41(16), 4630–4640. 10.1002/hbm.25146

Staudinger, M. R., Erk, S., & Walter, H. (2011). Dorsolateral prefrontal cortex modulates striatal reward encoding during reappraisal of reward anticipation. Cerebral Cortex, 21(11), 2578–2588.

van Driel, J., Sligte, I. G., Linders, J., Elport, D., & Cohen, M. X. (2015). Frequency band-specific electrical brain stimulation modulates cognitive control processes. PLoS One, 10(9), e0138984.

Vassena, E., Silvetti, M., Boehler, C. N., Achten, E., Fias, W., & Verguts, T. (2014). Overlapping neural systems represent cognitive effort and reward anticipation. PLoS One, 9(3), e91008. 10.1371/journal.pone.0091008

Vassiliadis, P., Beanato, E., Popa, T., Windel, F., Morishita, T., Neufeld, E., … Hummel, F. C. (2024). Non-invasive stimulation of the human striatum disrupts reinforcement learning of motor skills. Nature Human Behaviour, 8(8), 1581–1598. 10.1038/s41562-024-01901-z

Violante, I. R., Alania, K., Cassara, A. M., Neufeld, E., Acerbo, E., Carron, R., … Grossman, N. (2023). Non-invasive temporal interference electrical stimulation of the human hippocampus. Nat Neurosci, 26(11), 1994–2004. 10.1038/s41593-023-01456-8

Vriens, T., Vassena, E., Pezzulo, G., Baldassarre, G., & Silvetti, M. (2025). Meta-Reinforcement Learning reconciles surprise, value, and control in the anterior cingulate cortex. PLoS Comput Biol, 21(4), e1013025. 10.1371/journal.pcbi.1013025

Walton, M. E., Groves, J., Jennings, K. A., Croxson, P. L., Sharp, T., Rushworth, M. F., & Bannerman, D. M. (2009). Comparing the role of the anterior cingulate cortex and 6-hydroxydopamine nucleus accumbens lesions on operant effort-based decision making. European Journal of Neuroscience, 29(8), 1678–1691.

Webber, H. E., Lopez-Gamundi, P., Stamatovich, S. N., de Wit, H., & Wardle, M. C. (2020). Using pharmacological manipulations to study the role of dopamine in human reward functioning: A review of studies in healthy adults. Neuroscience & Biobehavioral Reviews.

Wessel, M. J., Beanato, E., Popa, T., Windel, F., Vassiliadis, P., Menoud, P., … Hummel, F. C. (2023). Noninvasive theta-burst stimulation of the human striatum enhances striatal activity and motor skill learning. Nat Neurosci, 26(11), 2005–2016. 10.1038/s41593-023-01457-7

Westbrook, A., Kester, D., & Braver, T. S. (2013). What is the subjective cost of cognitive effort? Load, trait, and aging effects revealed by economic preference. PLoS One, 8(7), e68210. 10.1371/journal.pone.0068210

Westbrook, A., Lamichhane, B., & Braver, T. S. (2019). The subjective value of cognitive effort is encoded by a domain-general valuation network. Journal of Neuroscience, 39(20), 3934–3947.

Westbrook, A., van den Bosch, R., Maatta, J. I., Hofmans, L., Papadopetraki, D., Cools, R., & Frank, M. J. (2020). Dopamine promotes cognitive effort by biasing the benefits versus costs of cognitive work. Science, 367(6484), 1362–1366. 10.1126/science.aaz5891

Wolpe, N., Holton, R., & Fletcher, P. C. (2024). What is mental effort: a clinical perspective. Biological Psychiatry, 95(11), 1030–1037.

Yohn, S. E., Errante, E. E., Rosenbloom-Snow, A., Somerville, M., Rowland, M., Tokarski, K., … Salamone, J. D. (2016). Blockade of uptake for dopamine, but not norepinephrine or 5-HT, increases selection of high effort instrumental activity: implications for treatment of effort-related motivational symptoms in psychopathology. Neuropharmacology, 109, 270–280.

Zalocusky, K. A., Ramakrishnan, C., Lerner, T. N., Davidson, T. J., Knutson, B., & Deisseroth, K. (2016). Nucleus accumbens D2R cells signal prior outcomes and control risky decision-making. Nature, 531(7596), 642–646. 10.1038/nature17400

